# Oxygen Sensor-Guided Fine Needle Biopsy Studies of Human Cancer Xenografts in Mice

**DOI:** 10.1101/2024.05.27.596060

**Authors:** Robert C. McDonald, Andrew H. Fischer, Mary Rusckowski

**Affiliations:** Giner, Inc. 89 Rumford Avenue Newton, MA 02466-1311; University of Massachusetts Medical School, Director, Cytopathology; University of Massachusetts Medical School, Associate Professor, Department of Radiology

**Keywords:** oxygen diffusion, hypoxia, breast carcinoma, adenocarcinoma, tumor xenograft in mice

## Abstract

An oxygen sensor-mounted fine-needle biopsy tool was used for *in vivo* measurement of oxygen levels in tumor xenografts. The system provides a means of measuring the oxygen content in harvested tumor tissue from specific locations. Oxygen in human tumor xenografts in a murine model was observed for over 1 min. Tissues were mapped in relation to oxygen tension (pO_2_) readings and sampled for conventional cytological examination. Careful modeling of the pO_2_ readings over 60 seconds yielded a diffusion coefficient for oxygen at the sensor tip, providing additional diagnostic information about the tissue before sampling. Oxygen level measurement may provide a useful adjunct to the use of biomarkers in tumor diagnosis.

## BACKGROUND

Fine-needle biopsy (FNB) is a well-established diagnostic method for minimally invasive sampling of cells and tissues when cancer is suspected. Over one million needle biopsies are performed annually in the United States alone^1^. FNB uses a 20-gauge or smaller needle inserted into the suspected regions of tissue for extraction of a tissue sample for microscopic analysis. In many cancers, the biopsied cells can be further analyzed using immunohistochemistry, molecular studies of nucleic acids, and analysis of a variety of emerging biomarkers to determine the type of cancer and its response to particular therapies^2^. Needle placement can be assisted by ultrasound, X-ray stereotactic, or magnetic resonance imaging to more precisely locate contrasted suspect tissue. For FNB samples to be adequate for diagnosis, it is crucial to obtain sufficient viable tumor cells. As we learn more about tumor biology, an adequate biopsy will also need to sample biologically heterogeneous parts of cancers to understand the mechanisms of action of new targeted therapies and to determine the optimal therapy for a particular patient. For most cancers, an FNB can be sufficient for basic diagnosis because the amount of tissue required for light microscopic diagnosis is relatively small, and ancillary immunohistochemistry and molecular testing require even smaller amounts of tissue or cells. The alternative to FNB is to use much larger biopsy devices, which cause considerably more discomfort and pose greater risks of hemorrhage/hematoma formation and infection. Efforts to improve FNB by allowing more representative sampling and avoiding sampling of necrotic or non-viable regions would spare many patients with the need for larger (core) biopsies. FNB with aspiration and front-end collection can harvest several tissue samples from different regions of the tumor without reinsertion of the device. The addition of an oxygen sensor located at the distal collecting end would synergize with the ability of micro sized devices to provide a sufficient biopsy sample to initiate appropriate therapies. In addition, minimally invasive FNB can permit samples to be obtained during therapy for established cancers to monitor the efficacy of treatment. It is impractical to sample cancers during therapy using larger biopsies.

In addition to tissue histology, pO_2_ measurements can provide diagnostic and immediate information about the nature of a suspicious lump or pathological process, independent of and synergistic with routine histopathologic diagnosis. Hypoxia is often associated with a worse patient prognosis^3^ This is especially true for hypoxic regions, where the temporal and spatial variation of oxygen can reveal information about the progress of the disease^4,5^. This study was conducted to demonstrate an oxygen-guided biopsy device with a fiber-optic phosphorescent oxygen sensor at the needle tip that permits rapid, real-time *in vivo* mapping of tissue oxygen levels. In addition to guiding better quality biopsies, this technology can potentially be used to assess disease progression, determine optimal therapies that are known to be dependent on tissue oxygen levels, and assess the efficacy of anti-angiogenic therapies^6–9^.

Vascular functional deficiency resulting in hypoxia has been studied in several cancers and is a recognized “Hallmark of Cancer”^10^. Currently used imaging methods are inadequate for locating regions with altered vasculature. Near-infrared (NIR) tomography has been used successfully to noninvasively image organs such as the breast, albeit at low resolution. Measurement of the NIR absorbance of oxygenated hemoglobin provides a means for imaging different levels of vasculature from elevated angiogenesis in malignant tissues^11^ is under development. However, this method is limited to superficial vasculature (1-2 mm depth) owing to light scattering and water absorption^12^. Magnetic resonance-guided near-infrared imaging (MRg-NIR) tissue levels of oxygen are indicative of differences in vascular compliance between normal and malignant breast tissue^13^. Blood oxygen-dependent MRI (BOLD-MRI) permits non-invasive mapping of normal and hypoxic vasculature, but only at low resolution^14^. While the BOLD-MRI and MRg-NIR methods were successful in distinguishing healthy tissue from abnormal tissue, it is unlikely that the methods can provide routine, rapid clinical results because of the complexity, size, and cost of the equipment, as well as the data processing time and incompatibility with metallic biopsy needles. Photoacoustic Imaging (PAI) has been demonstrated to be an effective means of high-resolution studies of tumor architecture using chemical probes, although the equipment may be out of reach for most clinical budgets and does not allow for continuous real-time monitoring of tumor oxygen level *in vivo*^15^.

The *ex vivo* MRI imaging to date points to the need for a more practical device for monitoring oxygen levels at depth with high spatial resolution in suspected cancer tissue *in vivo* to assess the relative levels of viable tumor, necrotic tumor, and normal cells present in a region of interest. A pO_2_ sensing biopsy tool would be an important adjunct to the collection of tissue samples, while gaining information on the metabolic state of a tumor^16^, and helping the development of hypoxia-related therapies. There are many potential applications; for example, hypoxia inducible factor 1 (HIF-1*α*) is overexpressed in tumor cells, promoting angiogenesis. HIF-1*α* is stabilized as oxygen levels decline^16^. Several groups are developing therapies based on the inhibition of HIF-1*α*. Targeted measurement of oxygen levels in hypoxic areas of tissue might provide a means of monitoring the efficacy of the treatment over time^17–19^. Conventional glass electrochemical sensors are unsuitable for minimally invasive needle probe devices because of their large diameters. Fiber-optic probes with much smaller diameters and excellent light transmission can be prepared. The probe used in this study was a fast-response phosphorescent O_2_-sensitive material mounted at the tip of an optical fiber close to the distal tip of the biopsy needle. The device and its development have been described elsewhere^20^.

This approach permits the biopsy of difficult cases presenting as small clusters of viable cancer cells within large necrotic areas that are expected to have different oxygen levels. For example, an overall average pO_2_ value below 15 mmHg was found for uterine, cervical, head and neck, and breast cancers compared to a normal value of ∼80 mmHg^21^. Tumor hypoxia is more pronounced in immunocompromised patients than in non-immunocompromised patients. Detection of hypoxia *in vivo* may be an important aid in identifying patients requiring more intensive treatment schedules^22^. Microbiopsies guided by pO_2_ could provide a means of sampling the highest, average, and lowest oxygenated regions to increase the understanding of cancer biology.

This paper describes a study undertaken in an animal model to evaluate an oxygen-sensing fiber optic probe incorporated in a fine-needle biopsy tool to 1) ensure sampling of viable (living) parts of a tumor, 2) provide clinicians with oxygen levels and spatial profiles to aid in choosing the most effective treatment therapies, and 3) aid in predicting clinical outcomes from selected therapies.

## METHODS

### Oxygen measurement protocol and Data treatment

Oxygen *in vivo* readings were recorded using a Microx 4™ Fiber Optic Oxygen Meter fitted with a PSt6 needle-type microsensor (Presens GmbH, Regensburg, Germany) to create an Oxygen-Guided Biopsy Needle (OGBN) as described in a separate publication^20^. The PreSens transmitter excites the phosphor at the fiber tip with a 505-nm light emitting diode and measures the resulting phosphorescence at 600-nm using a charge-coupled display device. The tool software was set to correct readings of subject body temperature and to report values every 1 second. Each reading is the average of five measurements. The oxygen-sensing optical fiber tip was 50 μm in diameter, within the oxygen diffusion limit of 100-150 μm in tissue^23^.

### Subject animal care

Female mice (Swiss National Institutes of Health) were procured from Charles River Laboratories. All *in vivo* measurements were conducted at the University of Massachusetts Small Animal Imaging Core Facility under a protocol approved by the University Animal Care and Use Committee (Animal Welfare Assurance (AWA) Number: A3306-01). Throughout the course of the study, animals showed no changes in appearance, inactivity, or signs of discomfort such as lethargy, shivering, immobility, hunched posture, weight loss, or ruffled fur. Tumors were not grown beyond 1.5 cm, in order to avoid interference with the animal’s mobility. When tumors reached the required size, the oxygen level was measured over 60 seconds following anesthesia, followed by fine-needle biopsy. The animal was then euthanized and the tumor was removed for histological analysis.

### Human breast cancer model

Tumors were initiated using breast carcinoma MDA-MB-231 (RRID:CVCL_0062) and adenocarcinoma LS-174T (RRID:CVCL_1384) cell line xenografts (ATTC, Manassas VA) in female nude mice. Primary tumors were initiated in the mammary pads of anesthetized female mice through injection of 4-5 × 10^6^ MDA-231 cells suspended in 0.05 mL media. Tumors were grown to three sizes (n=4):70-100 mm^3^ (approximately 14 days), 200-300 mm^3^ (21 days) and 400-600 mm^3^ (28 days) for the measurement of pO_2_ levels using the OGBN. With the selected number of injected cells and time intervals, tumors at 400-600 mm^3^ are predicted to develop intratumoral necrosis, an appropriate endpoint to meet the requirements of this investigation. After the tumor cells were injected into the mammary fat pad, the animals were observed to see that they were moving and behaving normally. They were checked again on days 4 and 7 for signs of tumor growth. Measurable tumors were present by day 7, after which they were measured with calipers every alternate day until they met the required size. During this time, the animals’ overall health was assessed.

During pO_2_ measurements, animals were set on a heating pad to maintain body temperature and anesthesia was maintained with 2-2.5% isoflurane (with oxygen and air supplied at a ratio of 0.2 L/min oxygen:0.8 L/min air) delivered by nose cone. At the required size, pO_2_ in tumors was measured at different locations to profile the complex heterostructures of tumor and non-tumor cells. During the procedure, the animals were maintained on a heating pad to maintain their body temperature. Following tumor biopsy aspiration, animals were euthanized and the extracted cells fixed for histological evaluation.

## RESULTS

### Tumor Measurements

Human MDA-MB-231 tumor growth was monitored using prototype OGBN probes with 25-Ga needles, equipped with a fiber-optic sensor. MDA-MB-231 orthotropic tumors were examined in mice at growth intervals of two and four weeks. Tumors grew in the left and right mammary fat pads. **Figure 1** shows the bench setup with the animal positioned for the OGBN measurements with a micropositioner to control the needle location. The temperature correction for the phosphor used for the PreSens optical fiber sensor was set to 37°C °C using the Microx 4 meter.

**Figure 1.**
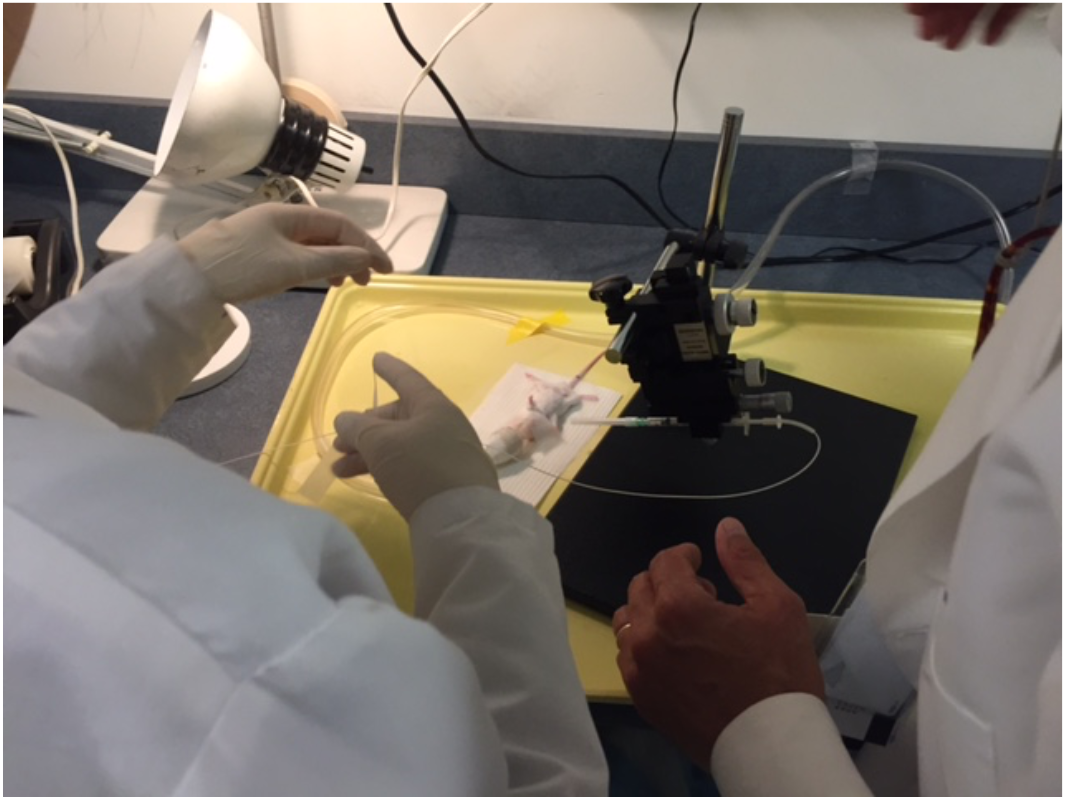
Bench setup with animal positioned for OGBN measurements

Orthotropic tumors examined *in vivo* showed hypoxic regions starting in the second week with decreasing levels of oxygen over 4 weeks for both right and left mammary tumors. Oxygen levels showed a complex relationship with tumor necrosis. Human LS-174T colon cancer xenografts in flanks presented with an exceptionally homogenized mixture of necrotic and viable tumor cells (Figure 2), making them suitable for determining the average pO_2_ for decay curve models (see below). The sample is shown again in Figure 8 for comparison with the mostly necrotic cells.

**Figure 2.**
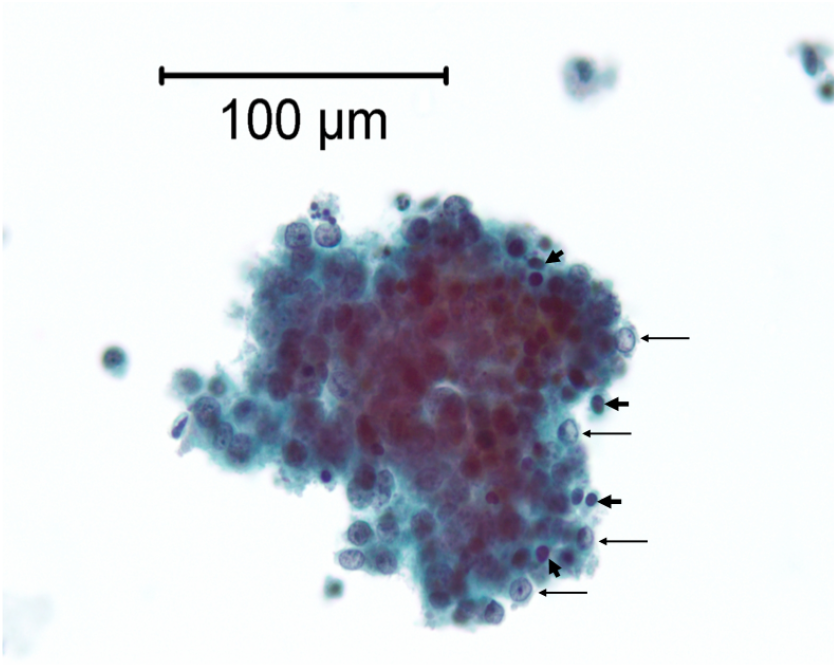
Human colon cancer xenograft, obtained using an oxygen sensor-coupled Fine Needle Aspirated (FNA) needle used for these studies, displayed in a non-embedded, Papanicolaou-stained ThinPrep slide. There is an intimate admixture of viable cells with euchromatic nuclei (some shown with long-thin arrows) and dead cells (a few shown with bold arrowheads) as indicated by their smaller darker pycnotic nuclei.

Figure 3 shows one of the MDA-MB-231 tumors as a montage of tissue photomicrographs following *in vivo* measured % oxygen saturation at 1-mm intervals (1 mm/60 seconds). The location of the needle was mapped by tattooing the entrance and exit sites with different colors, excising and fixing the tumor, and then performing serial histological sections through a plane that included the entrance and exit sites. Tissue compression was corrected during the microtome sectioning. Tissue shrinkage was not observed during fixation or embedding.

**Figure 3.**
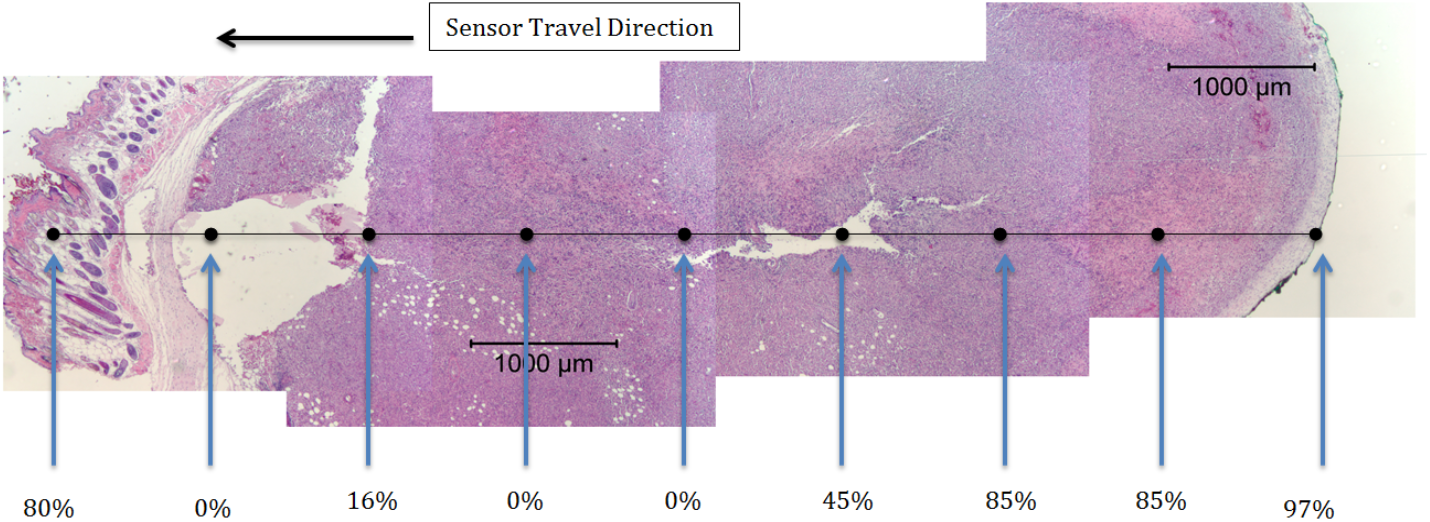
Percent O_2_ saturation in 4-week-old MDA-MB-231 xenograft; white areas represent needle-torn tissue

The entry and exit locations for the OGBN linear measurements in the tumors were marked with indelible ink so that the excised tumor could be oriented for sectioning, embedding, and optical microscopy. Very hypoxic (0% O_2_ saturation) regions were clearly observed with the device, although a strict correlation between necrosis and oxygen measurements in MDA-MB-231 xenografts was not always observed. For example, the second measurement (85%) mapped to an area with coagulative necrosis, while the third measurement (85%) occurred in an ostensibly viable area, and the 6^th^ measurement (0%) also appeared to be viable based on the histologic appearance. Although Figure 3 represents only a single section, it is possible that thin sections above and below this plane would result in a stricter correlation between histology and oxygen levels.

A second xenograft (LS-174T colonic adenocarcinoma) was examined in mice because of its faster growth and lower extent of necrosis. Nine tissue samples were extracted using the OGBN developed with a side port to permit the removal of each tissue sample with a saline rinse. This prevents residual pO2 from tissue carryover. Saline rinse can eliminate oxygen carryover between tissue samples. Each sample was removed after a 60-seconds recording of pO2 levels as oxygen diffused to the sensing phosphor. All LS-174T tumor samples showed a relatively homogeneous mixture of necrotic and viable cancer cells in this xenograft with a diameter of 10-15 microns (Figure 2 above). Figure 4 and Figure 5 show the recorded pO_2_ values for nine FNA biopsies obtained with the OBGN probe procured from the left and right flank colon xenograft tumors of one animal. Fine-needle aspirated (FNA) samples were prepared using a ThinPrep® (Hologic, Inc.) technique that deposits a thin layer of suspended tissue fragments in a 20-mm<u>-diamete</u>r circula<u>r area.</u> The cell yield was very high. ThinPrep slides prepared from mostly necrotic cells (Test 2, light blue) and primarily viable tumor cells (Test 4, green) are shown in Figure 6. The measured oxygen levels were low for most necrotic tissue samples.

**Figure 4.**
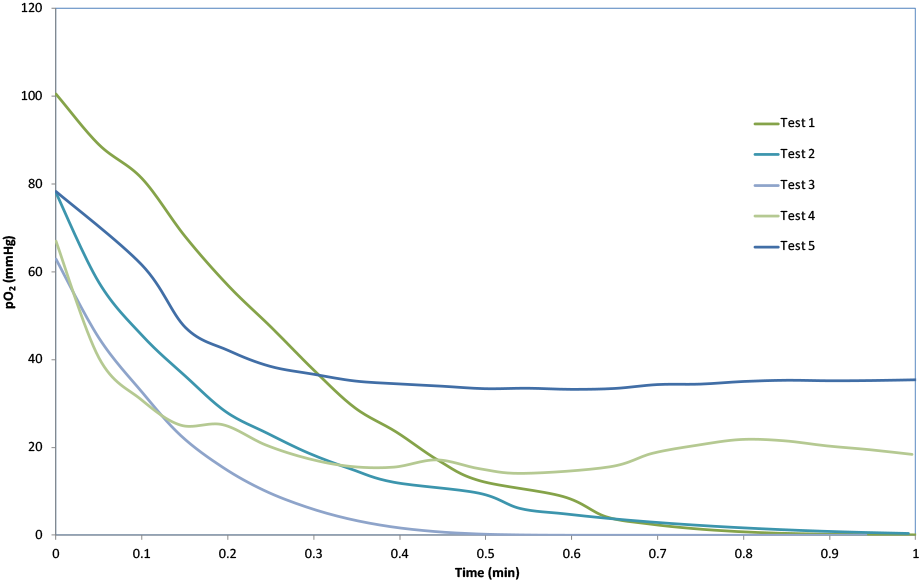
Tissue oxygen measured over 1 minute prior to tissue extraction from left flank LS-174T xenograft

**Figure 5.**
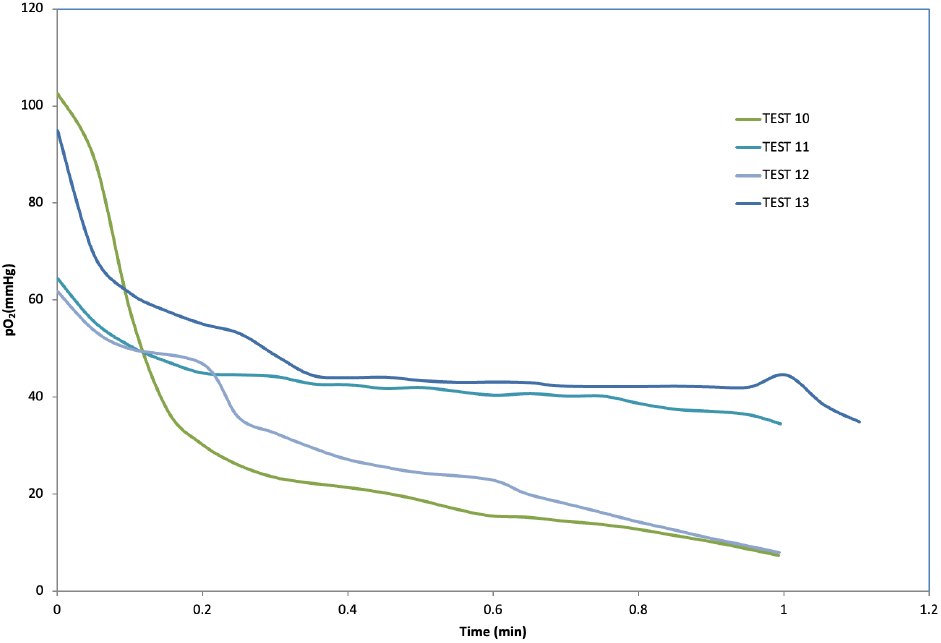
Tissue oxygen measured over 1 minute prior to tissue extraction from right flank LS-174T xenograft

**Figure 6.**
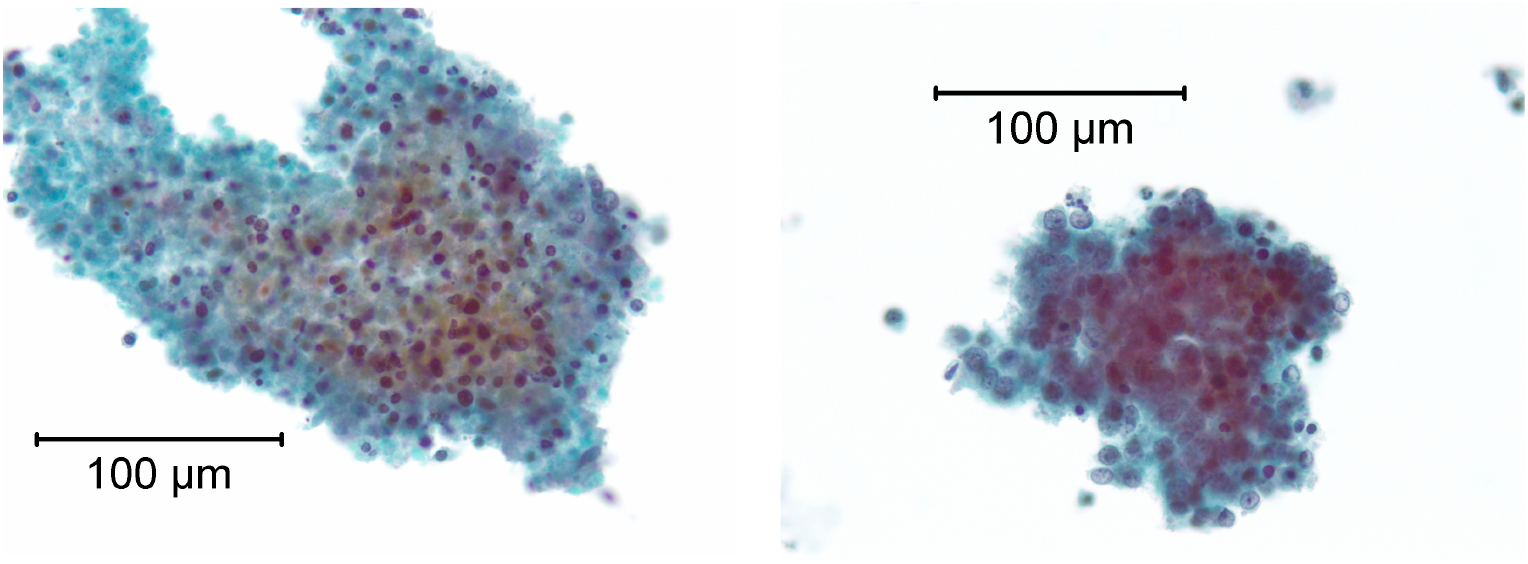
Xenograft colon tumor samples obtained by an oxygen sensor-coupled FNA (fine needle aspiration) needle from a left flank as shown in ThinPrep slides, Papanicolaou-stained at 400X magnification. On the left is a mostly necrotic sample (Test 2), with only small pycnotic nuclei or no residual nuclei. On the right is a mostly viable tumor sample (Test 4).

**Commented [P1]: Conventions:** Abbreviations are usually defined at the first use in the abstract as well as in the main text. Check whether ‘FNA’ should be defined here.

### Oxygen diffusion model for tumor measurements

During the placement of the biopsy tool, just prior to taking a tissue sample, the sensor shows a declining pO_2_ from the initial value in air until the ambient tissue concentration of O_2_ is established at the phosphor surface. This process may provide additional information regarding tissue samples related to cell physiology and metabolism. Each measurement started high owing to the presence of atmospheric oxygen entering the sensor between measurements when the sensor was exposed to

air. No oxygen is consumed with this type of oxygen quenching phosphor, so the decline in pO_2_ is the result of oxygen diffusion from the sensor to the surrounding tissue until equilibrium is reached. The pO_2_ decay curve can be fitted to a double exponential function by immediate placement of the sensor at a given location:

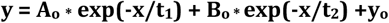

where y is the pO_2_ reading at a fixed location over time and x is the time in seconds. *t*_*1*_ and *t*_*2*_ represent the oxygen diffusion constants of the sensor and tissue, respectively. *A*_*o*_ and *B*_*o*_ are the relative amplitudes of the two regions during the overall diffusion process. Six measurements were taken in one adrenal carcinoma xenograft approximately 150-230 microns in size. PO_2_ readings were recorded at 3-second intervals, while the sensor was held close to the sampling distal end of the needle, in contact with the tissue. Tests 8 and 6 in Figure 6 show representative traces of PO_2_ measured in viable tumor (18-38 mmHg) and hypoxic tissue (<1 mmHg), respectively. The observed equilibrium oxygen concentration and diffusion parameter fits for each tissue sample are listed in Table 1.

**Table 1.** Double-exponent fits for 1-minute decay in measured pO_2_ in colon tumor xenografts.

| | Lowest $pO_2$<br>(mmHg) | $y_o$ | $A_o/B_o$ | $A_o$ | $t_1$ | $B_o$ | $t_2$ | Adj. $R^2$ |
| --- | --- | --- | --- | --- | --- | --- | --- | --- |
| Test 1 | 1.21 | -7.38 | 0.930878 | 55 | 0.3163 | 59 | 0.32 | 0.9914 |
| Test 2* | 0.37 | -0.84 | 1.140515 | 31 | 0.25455 | 27 | 0.25 | 0.9884 |
| Test 3 | 0.00 | -1.91 | 0.991918 | 33 | 0.14688 | 33 | 0.15 | 0.9998 |
| Test 4* | 14.70 | 17.96 | 0.118076 | 5 | 0.00124 | 44 | 0.08 | 0.9519 |
| Test 5 | 35.10 | 33.72 | 0.880387 | 22 | 0.1285 | 25 | 0.13 | 0.9608 |
| Test 10 | 4.50 | -791004 | 0.000109 | 86 | 0.11412 | 791026 | 61950. | 0.9735 |
| Test 11 | 28.20 | -3.63E+06 | 4.84E-06 | 18 | 0.07083 | 3625730 | 331128. | 0.9888 |
| Test 12 | 8.00 | -1.07E+06 | 2.58E-05 | 28 | 0.24231 | 1070370 | 41087. | 0.9838 |
| Test 13 | 42.10 | 40.50 | 0.55682 | 19 | 0.01825 | 35 | 0.21 | 0.97404 |

\*Stained tissue micrographs are shown in Figure 6.

The PSt6 low-oxygen sensors used in this study had a diameter of 140 μm. This dimension is too large to detect differences in the necrotic and viable tumor regions shown in Figure 6, but small enough to yield an average of pO_2_ values over the single tissue sample.

The data can be interpreted in terms of the relative amplitudes *Ao* for oxygen in the phosphor/binder complex ranging from 5 to 86 mm Hg/min. and *Bo* for oxygen in the surrounding tissues, reflecting the degree of hypoxia. The values for amplitude *Bo* in Tests 10-12 of the right-side tumor samples were exceptionally high, indicating that the surrounding tissue dominated the sensor response to a much greater extent. The ratio *Ao/Bo* provides an estimate of the relative diffusion limitations of the sensor itself compared to the surrounding tissue. The table also lists the calculated pO_2_ intercept Y_o_ and the lowest measured pO_2_ over one minute. The two columns agree well for most measurements, supporting the double-exponent model. The unreasonably low calculated intercepts, Y_o_, for these lowest calculated pO_2_ values suggest that a more complex model is necessary.

Overall, oxygen measurements in mouse xenograft tumors showed variations in pO_2_ at 1- to 2-mm spatial resolution. The technique offers a high degree of detail on tumor structure and the distribution of tumor, non-tumor, and necrotic tissues typical of heterogeneous breast cancers. The human breast tumor xenograft does not show a clear mapping of oxygen levels suspected for necrotic and viable tumor cells, which will require larger tumor models for clarification. Several experiments were conducted using the two xenografts in mice to explore the potential use of the OGBN tool using both 25-Ga and 21-Ga needles. We found:

- The oxygen probe was placed at a range of locations within or slightly beyond the opening of the needle without affecting the pO_2_ reading.
- Viable tissues could be sampled without disturbing the phosphor over several insertions.

The pO_2_ response time was significantly longer in the less hypoxic regions as shown in Figure 7, generated from a series of different tumor regions measured at depths of 2-6 mm. The double exponential fit, using an orthogonal distance regression routine, is consistent with at least two different oxygen diffusion rates, first in tumor tissue and second in the silicone-bonded oxygen-sensing phosphor itself.

**Figure 7.**
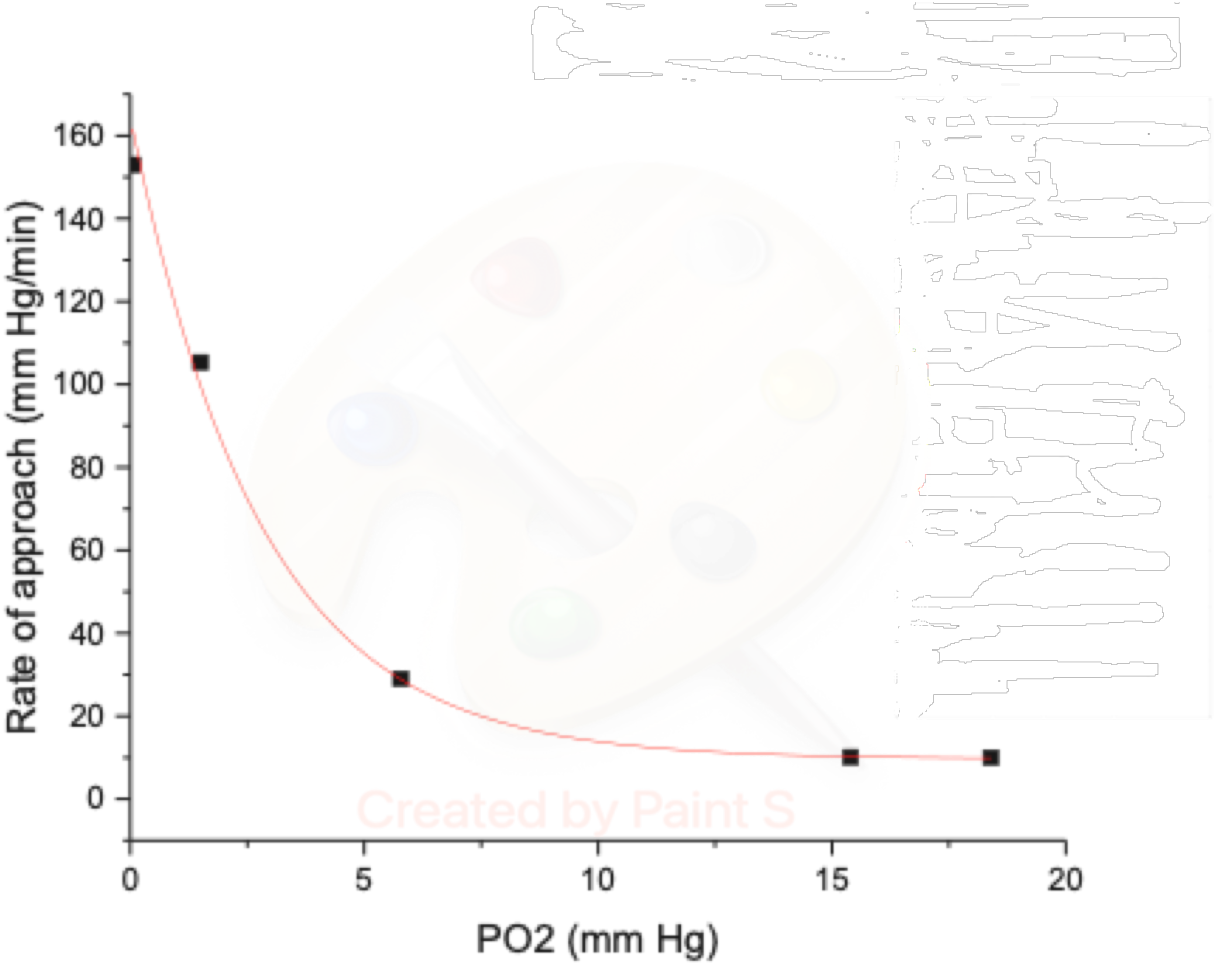
Showing complex, double exponential relationship between pO_2_ and the rate at which sensor reaches steady state

## DISCUSSION

The rapid response of the OGBN device in tissue results from the normal scavenging of singlet oxygen species formed at the phosphor by non-enzymatic molecules, such as glutathione and antioxidant enzymes present in the intercellular region^24^. Thus, the falloff in pO_2_ readings is not due to the transient buildup of singlet oxygen but rather to the slow fluctuation of molecular oxygen. Variable oxygen levels are not unexpected^25^. *In vivo* results with oral and nasal canine tumors obtained with optical (OxyLite™) probes showed evidence of pO_2_ fluctuations in hypoxic tumors with amplitudes of 2-20 mmHg and periods of 14–40 minutes^26^. However, these measurements did not permit the precise location of the probes or accurate measurements during the first 10 minutes of insertion. Also, the device did not provide the ability to remove tissue at each site and correlate histology with oxygen levels.

This work validated the value of using an OGBN to continuously read pO_2_ at the site of a fine-needle biopsy to examine temporal changes at key locations in tumors. 20 Seconds of steady state transport of O_2_ diffusion in the sensing phosphor provides a reliable record of pO_2_ values and fluctuations. Necrotic, non-informing cells located > 200 micons from a blood vessel will exhibit extreme hypoxia^21^ permitting the pathologist to avoid sampling.

Recent reports suggest that tumor oxygen levels can be coupled with a second biomarker such as hypoxia inducible factor (HIF) for certain tumors to distinguish necrotic from viable tumor tissue^27^. While the extent of hypoxia can be estimated from the presence of biomarkers such as HIF-1*α*, osteopontin, and vascular endothelial growth factor (VEGF), detailed oxygen levels in different regions of the tumor are lacking^28,29^. The OGBN probe can thus confirm hypoxia while permitting more detailed mapping. Oxygen levels are important in determining the efficacy and appropriateness of radiotherapy (RT)^30,31^. Hypoxic cells are especially resistant to RT, with oxygen levels below 20 mmHg and rapid falloff in effectiveness at 3 mmHg^29,31^. The tumor microenvironment can become resistant to normal T-cell immune response, as hypoxia drives elevated levels of and concomitant suppression of A2 adenosine receptors on the surface of activated immune cells^32^.

The distribution of oxygen levels in tumors are complex^33^ and as yet poorly understood. Improved techniques, such as the present one, will aid in measuring *in vivo* levels accurately and in real time. The FNA pO_2_ probe permits rapid measurement of oxygen levels in tumors at precise locations. The measured oxygen levels in MDA-231 implanted tumors in mice exhibited cycling hypoxia over 1-5 mins. One possibility is that after a region of the tumor has undergone ischemic necrosis, there is no longer a drain on pO_2_ from the metabolism of viable tumor cells. Oxygen diffusion into the necrotic area can occur over time. Changes in oxygen levels within *in vivo* tumor xenografts also show evidence of autofluorescence associated with cancer-related metabolism in epithelial cancer stem cells in multiple types of human tumors ^34^. This technique may be useful for determining tumor staging. Recent findings indicate that low-oxygen stem cells in the intercellular matrix add additional information about the extent of metastasis^3534^.

## CONCLUSION

A method was developed so that linear traces of pO_2_ at 1- to 2-mm intervals could be overlaid onto histological sections fixed from excised tumors. Slowly varying pO_2_ levels reflecting physiological changes in tumors require further study. This approach permits the biopsy of difficult cases presenting as small clusters of viable cancer cells within large necrotic areas that are expected to have different oxygen levels. For example, an overall average pO_2_ value of less than 20 mmHg was found for uterine, cervical, head and neck, and breast cancers compared to a normal value of 80 mmHg. Tumor hypoxia is more pronounced in immunocompromised patients, and the detection of hypoxia *in vivo* can be an important aid in identifying patients requiring more intensive treatment schedules^37^. Microbiopsies guided by pO_2_ could provide a means of sampling the highest, average, and lowest oxygenated regions to increase the understanding of cancer staging. Given the likelihood of oxygen diffusion from viable tissue into and through necrotic tissue, the measurement of elevated oxygen may be the result of diffusion into necrotic tissue from the surrounding

vasculature. Thus, the full effectiveness of pO_2_ readings requires supplemental histology of aspirated cells and/or molecular biomarkers.

## Author Information

***Authors***

**Andrew H. Fischer -** *Department of Pathology, University of Massachusetts Medical School, 55 Lake Avenue, Worcester, Massachusetts 01655, Orcid No.*

**Mary Rusckowski -** *Department of Radiology, University of Massachusetts Medical School, 55 Lake Avenue, Worcester, Massachusetts 01655, Orcid No.*

## AUTHOR CONTRIBUTIONS

**R**.**C. McDonald**: Conceptualization, data curation, formal analysis, investigation, visualization, methodology, writing–original draft, supervision, funding acquisition, project administration.

**A**.**H. Fischer**: Formal tissue analysis, investigation, resources.

**M. Rusckowski:** Resources, equipment, animal curation.

## Acknowledgements

The authors thank Michael Riera-Smith for assisting with probe characterization and Yuzhen Wang for animal care. This work was funded by the National Institutes of Health, National Cancer Institute under Contract No. HHSN261201400056C.

## References

1. John M. Eisenberg Center for Clinical Decisions and Communications Science. Core-Needle Biopsy for Breast Abnormalities. In: Comparative Effectiveness Review Summary Guides for Clinicians. AHRQ Comparative Effectiveness Reviews. Agency for Healthcare Research and Quality (US); 2007. Accessed June 9, 2021. http://www.ncbi.nlm.nih.gov/books/NBK368367/

2. Jiang L, Lin X, Chen F, et al. Current research status of tumor cell biomarker detection. Microsyst Nanoeng. 2023;9(1):123. doi:10.1038/s41378-023-00581-5

3. Godet I, Doctorman S, Wu F, Gilkes DM. Detection of Hypoxia in Cancer Models: Significance, Challenges, and Advances. Cells. 2022;11(4):686. doi:10.3390/cells11040686

4. Huang J, Shen J, Qiu Y, et al. Abstract 5002: The prognostic relevance and underlying mechanisms of the novel oxygen sensor ADO in cancers. Cancer Res. 2022;82(12_Supplement):5002–5002. doi:10.1158/1538-7445.AM2022-5002

5. Li Y, Zhao L, Li XF. Hypoxia and the Tumor Microenvironment. Technol Cancer Res Treat. 2021;20:153303382110363. doi:10.1177/15330338211036304

6. Jászai J, Schmidt M. Trends and Challenges in Tumor Anti-Angiogenic Therapies. Cells. 2019;8(9):1102. doi:10.3390/cells8091102

7. McIntyre A, Harris AL. Metabolic and hypoxic adaptation to anti-angiogenic therapy: a target for induced essentiality. EMBO Mol Med. 2015;7(4):368–379. doi:10.15252/emmm.201404271

8. Rapisarda A, Melillo G. Role of the hypoxic tumor microenvironment in the resistance to anti-angiogenic therapies. Drug Resist Updat. 2009;12(3):74–80. doi:10.1016/j.drup.2009.03.002

9. Ulivi P, Marisi G, Passardi A. Relationship between hypoxia and response to antiangiogenic therapy in metastatic colorectal cancer. Oncotarget. 2016;7(29):46678–46691. doi:10.18632/oncotarget.8712

10. Hanahan D, Weinberg RA. Hallmarks of Cancer: The Next Generation. Cell. 2011;144(5):646–674. doi:10.1016/j.cell.2011.02.013

11. Cheung A, Tu L, Macnab A, Kwon BK, Shadgan B. Detection of hypoxia by near-infrared spectroscopy and pulse oximetry: a comparative study. J Biomed Opt. 2022;27(07). doi:10.1117/1.JBO.27.7.077001

12. Simpson CR, Kohl M, Essenpreis M, Cope M. Near-infrared optical properties of ex vivo human skin and subcutaneous tissues measured using the Monte Carlo inversion technique. Phys Med Biol. 1998;43(9):2465–2478. doi:10.1088/0031-9155/43/9/003

13. Carpenter CM, Rakow-Penner R, Jiang S, et al. Inspired gas-induced vascular change in tumors with magnetic-resonance-guided near-infrared imaging: human breast pilot study. J Biomed Opt. 2010;15(3):036026. doi:10.1117/1.3430729

14. Gloviczki ML, Saad A, Textor SC. Blood oxygen level-dependent (BOLD) MRI analysis in atherosclerotic renal artery stenosis: Curr Opin Nephrol Hypertens. 2013;22(5):519–524. doi:10.1097/MNH.0b013e32836400b2

15. Hui X, Malik MOA, Pramanik M. Looking deep inside tissue with photoacoustic molecular probes: a review. J Biomed Opt. 2022;27(07). doi:10.1117/1.JBO.27.7.070901

16. Paredes F, Williams HC, San Martin A. Metabolic adaptation in hypoxia and cancer. Cancer Lett. 2021;502:133–142. doi:10.1016/j.canlet.2020.12.020

17. Brown JM, Wilson WR. Exploiting tumour hypoxia in cancer treatment. Nat Rev Cancer. 2004;4(6):437–447. doi:10.1038/nrc1367

18. Semenza GL. Heritable disorders of oxygen sensing. Am J Med Genet A. 2021;185(8):2576–2581. doi:10.1002/ajmg.a.62250

19. Melillo G. Targeting hypoxia cell signaling for cancer therapy. Cancer Metastasis Rev. 2007;26(2):341–352. doi:10.1007/s10555-007-9059-x

20. McDonald RC. Development of a pO2-Guided Fine Needle Tumor Biopsy Device. J Med Devices. 2022;16(2):021003. doi:10.1115/1.4052900

21. McKeown SR. Defining normoxia, physoxia and hypoxia in tumours— implications for treatment response. Br J Radiol. 2014;87(1035):20130676. doi:10.1259/bjr.20130676

22. Vaupel P, Höckel M, Mayer A. Detection and Characterization of Tumor Hypoxia Using pO 2 Histography. Antioxid Redox Signal. 2007;9(8):1221–1236. doi:10.1089/ars.2007.1628

23. Eales KL, Hollinshead KER, Tennant DA. Hypoxia and metabolic adaptation of cancer cells. Oncogenesis. 2016;5(1):e190–e190. doi:10.1038/oncsis.2015.50

24. Reczek CR, Chandel NS. The Two Faces of Reactive Oxygen Species in Cancer. Annu Rev Cancer Biol. 2017;1(1):79–98. doi:10.1146/annurev-cancerbio-041916-065808

25. Gonçalves MR, Johnson SP, Ramasawmy R, Pedley RB, Lythgoe MF, Walker-Samuel S. Decomposition of spontaneous fluctuations in tumour oxygenation using BOLD MRI and independent component analysis. Br J Cancer. 2015;113(8):1168–1177. doi:10.1038/bjc.2015.270

26. Brurberg KG, Skogmo HK, Graff BA, Olsen DR, Rofstad EK. Fluctuations in pO2 in poorly and well-oxygenated spontaneous canine tumors before and during fractionated radiation therapy. Radiother Oncol. 2005;77(2):220–226. doi:10.1016/j.radonc.2005.09.009

27. Loo JM, Scherl A, Nguyen A, et al. Extracellular Metabolic Energetics Can Promote Cancer Progression. Cell. 2015;160(3):393–406. doi:10.1016/j.cell.2014.12.018

28. Vergis R, Corbishley CM, Norman AR, et al. Intrinsic markers of tumour hypoxia and angiogenesis in localised prostate cancer and outcome of radical treatment: a retrospective analysis of two randomised radiotherapy trials and one surgical cohort study. Lancet Oncol. 2008;9(4):342–351. doi:10.1016/S1470-2045(08)70076-7

29. Walsh JC, Lebedev A, Aten E, Madsen K, Marciano L, Kolb HC. The Clinical Importance of Assessing Tumor Hypoxia: Relationship of Tumor Hypoxia to Prognosis and Therapeutic Opportunities. Antioxid Redox Signal. 2014;21(10):1516–1554. doi:10.1089/ars.2013.5378

30. Lawrence YR, Dicker AP. Hypoxia in prostate cancer: observation to intervention. Lancet Oncol. 2008;9(4):308–309. doi:10.1016/S1470-2045(08)70081-0

31. Sørensen BS, Horsman MR. Tumor Hypoxia: Impact on Radiation Therapy and Molecular Pathways. Front Oncol. 2020;10:562. doi:10.3389/fonc.2020.00562

32. Sitkovsky M, Ohta A. Targeting the hypoxia-adenosinergic signaling pathway to improve the adoptive immunotherapy of cancer. J Mol Med. 2013;91(2):147–155. doi:10.1007/s00109-013-1001-9

33. Lopez-Serra P, Marcilla M, Villanueva A, et al. A DERL3-associated defect in the degradation of SLC2A1 mediates the Warburg effect. Nat Commun. 2014;5(1):3608. doi:10.1038/ncomms4608

34. Miranda-Lorenzo I, Dorado J, Lonardo E, et al. Intracellular autofluorescence: a biomarker for epithelial cancer stem cells. Nat Methods. 2014;11(11):1161–1169. doi:10.1038/nmeth.3112

35. Shi X, Zhang Y, Zheng J, Pan J. Reactive Oxygen Species in Cancer Stem Cells. Antioxid Redox Signal. 2012;16(11):1215–1228. doi:10.1089/ars.2012.4529

36. Dalerba P, Dylla SJ, Park IK, et al. Phenotypic characterization of human colorectal cancer stem cells. Proc Natl Acad Sci. 2007;104(24):10158–10163. doi:10.1073/pnas.0703478104

37. Vaupel P, Schlenger K, Knoop C, Ha M. Oxygen Distribution In Human Tumors: Evaluation of Tissue Oxygen Distribution in Breast Cancers by Computerized O2 Tension Measurements’. Cancer Res. 51:3316–3322. https://cancerres.aacrjournals.org/content/canres/51/12/3316.full.pdf

